# Locomotor savings relies on attentional control of walking in older, but not younger adults

**DOI:** 10.64898/2026.01.12.698834

**Authors:** Shuqi Liu, Andrea L. Rosso, Emma M. Baillargeon, Andrea M Weinstein, Gelsy Torres-Oviedo

## Abstract

The ability to recall learned movements and rapidly adapt to environmental changes, known as locomotor savings, is crucial for mobility in community-dwelling older adults. However, the influence of aging on locomotor savings and the underlying mechanisms remains poorly understood. Attentional compensation is a particularly relevant mechanism because the control of automatic motor behaviors like walking tend to recruit more attentional/executive resources with aging. We hypothesize that locomotor savings is diminished with age and relies on attentional rather than automatic control of walking. To test this, we compared savings of a novel walking pattern learned on a split-belt treadmill, where each leg moves at a different speed, across multiple days in 21 older and 21 younger adults. Attentional control of walking was assessed by overground dual-task walking while prefrontal cortex (PFC) activity was recorded using functional near-infrared spectroscopy (fNIRS). We found that older adults exhibited less locomotor savings than younger adults after practice. Older adults also relied more on attentional resources during dual-task walking. Importantly, greater locomotor savings was associated with higher attentional control of walking in older adults, suggesting that the use of attentional resources during challenging walking facilitates the recall of previously learned movements. These results indicate that cognitive compensation strategies utilizing attentional resources are important neural mechanisms modulating locomotor savings. Understanding the role of cognitive compensation in locomotor savings may inform rehabilitation design to enhance mobility in older adults ensuring movement corrections practiced in clinical settings are saved for long-term benefit in daily life.

## Introduction

Mobility in the community occurs within a dynamic environment and often requires adapting walking patterns according to the situations at hand and to recall previously adapted movements (1, 2), known as locomotor savings (3–6). For example, navigating icy surfaces initially requires reactive motor commands; however, subsequent encounters allow for the anticipatory retrieval of learned patterns, ensuring stability. Locomotor savings over multiple days is especially important in community mobility, where individuals must repeatedly draw on previously learned walking patterns to handle recurring challenges (1, 2, 7, 8), such as crossing busy intersections or navigating construction zones.

Inability to recall previously learned walking patterns when needed, i.e. poor locomotor savings, could contribute to restrictions in community-mobility, the ability to move outside one’s home (9), which are common (10, 11) in older ages. However, the effect of aging on multi-day locomotor savings remains unclear. Prior studies (12–14) suggested that older adults exhibited impaired locomotor savings when tested across days, but these findings are inconclusive due to uncontrolled intervals between testing days and the absence of practice to induce robust locomotor savings. In this study, we systematically test whether older adults can demonstrate locomotor savings across multiple days following a structured practice and quantify the extent of locomotor savings relative to younger adults.

Furthermore, we are interested in elucidating the neural mechanisms of locomotor savings, particularly the role of attentional compensation. Attention will be defined as the information processing capacity of the prefrontal cortex (PFC) here (15). With older age, walking control shifts from predominantly automatic to compensatory strategies, relying more heavily on attentional and executive resources (16). This shift in control is often assessed by dual-task walking (walking while performing an attention demanding task) with recording of prefrontal cortex (PFC) activity. In this context, decreased task performance and/or increased PFC activity indicate that walking is competing with the secondary task for attentional resources, which is interpreted as reduced automaticity and increased attentional control of walking (17–21). Consistently, reaching motor savings in younger adults was related to brain activation in the cortico-striatal loops (22–24) but in older adults was linked to more frontal brain activities (25). Locomotor savings in younger adults are typically linked to subcortical circuits, such as the cerebellar-thalamic-cortical and cerebellar-basal ganglia circuits (26), and these structures often undergo age-related atrophy (27, 28). Therefore, we hypothesize attentional compensation during walking will be relevant to locomotor savings in older adults.

In summary, this study investigates age-related differences in locomotor savings across multiple days and evaluates the contribution of attentional compensation. We hypothesize that with aging, locomotor savings is diminished and relies on attentional rather than automatic control of walking. To test this, we assessed locomotor savings and attentional control of walking in older and younger adults. Locomotor savings was quantified by retrieval of a novel walking pattern acquired through repeated exposure to a split-belt treadmill, where each leg moves at a different speed. Attentional control of walking was evaluated using a well-established overground dual-task walking paradigm (20, 29), with neural compensation from the PFC measured by functional near-infrared spectroscopy (fNIRS). Finally, to explore the clinical relevance of these findings in older adults, we tested the relationship between locomotor savings, dynamic balance and mobility as measured by the Functional Gait Assessment (FGA) (30), and cognitive abilities evaluated by neuropsychological assessment in older adults.

## Results

We recruited 21 older (>= 65 years old) and 21 younger (19-40 years old) adult participants in this study. Demographics and self-selected overground walking speeds are shown in Table 1. Leg dominance was determined by self-reported leg used to kick a soccer ball. Self-selected overground walking speed was calculated as the average speed walking back and forth on a straight 9.2m walkway for 150 strides (Figure 1A baseline overground). Older adults were screened for no signs of cognitive impairments as indicated by a Montreal Cognitive Assessment (MoCA) score greater than or equal to 24. The full inclusion criteria for both age groups are included in the Methods. The Functional Gait Assessment (FGA) was administered to older adults to evaluate dynamic balance.

**Table 1.** Participant characteristics.

| **Variables** | **Older Adults**<br>**Mean (SD)**<br>**or n (%)** | **Younger Adults**<br>**Mean (SD)**<br>**or n (%)** |
| --- | --- | --- |
| Sample size | 21 | 21 |
| Age (years) | 71.85 ± 3.14 | 25.05 ± 5.20 |
| Female, n (%) | 11 (52.38%) | 11 (52.38%) |
| Race, n (%) |  |  |
| White | 19 (90.48%) | 16 (76.19%) |
| Black or African American | 1 (4.76%) | 1 (4.76%) |
| More than one race | 1 (4.76%) | 0 |
| Asian | 0 | 4 (19.05%) |
| Highest level of education, n (%) |  |  |
| Secondary level education / GED / Equivalent | 2 (9.52%) | 0 |
| Some college | 4 (19.05%) | 8 (38.10%) |
| Bachelor's degree or higher | 15 (71.43%) | 13 (61.90%) |
| Right leg dominant, n (%) | 21 (100%) | 20 (95.24%) |
| Self-selected overground walking speed (m/s) | 0.96 ± 0.17 | 1.01 ± 0.14 |
| Montreal Cognitive Assessment (MoCA) score (max 30) | 27.05 ± 1.47 | not administered |
| Functional Gait Assessment (FGA) score (max 30) | 24.71 ± 2.43 | not administered |

**Figure 1.**
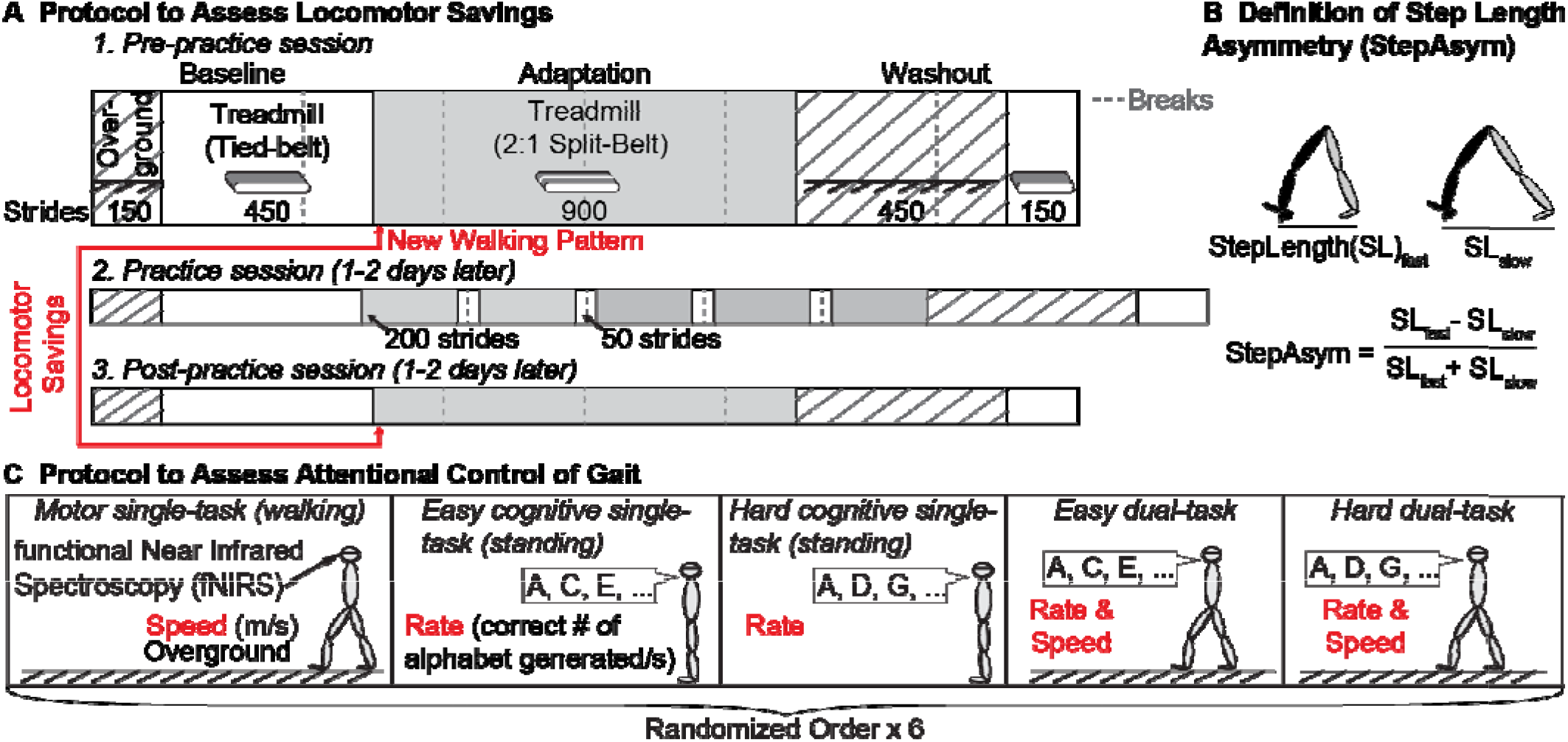
Experimental protocols and key outcome measures. (A) Schematics of the split-belt treadmill walking paradigm to evaluate locomotor savings. Participants completed three sessions on separate days: pre-practice, practice, and post-practice with each session 1 or 2 days apart. In the pre-practice session, participants learned a new walking pattern on a split-belt treadmill. Locomotor savings quantifies the ability to recall this newly learned pattern after practice and was measured by the change in step length asymmetry (StepAsym, B) in the post-compared to pre-practice session. Patched blocks represent self-paced overground walking, white blocks represent tied-belt treadmill walking, grey blocks represent split-belt walking, and dashed grey vertical lines represent breaks. (B) Illustrations of StepAsym parameter calculation where zero represents symmetric walking. Step length (SL) is defined as the distance between two ankles during heel strike. (C) Schematics of the dual-task walking paradigm with prefrontal cortical (PFC) recordings from functional near infrared spectroscopy (fNIRS) to evaluate attentional control of walking. Both motor (walking speed) and cognitive (correct number of alphabet letters generated per second) performances were measured. Participants performed motor (walking) and cognitive (reciting every two or three letters of the alphabet) tasks either separately (single-task) or simultaneously (dual-task) in randomized order for 6 repetitions.

### Older adults showed less locomotor savings than younger adults

Locomotor savings was evaluated by the change in step length asymmetry (StepAsym, Fig 1B) before and after completing a split-belt treadmill walking practice session (Figure 1A). StepAsym is defined as the normalized difference in bilateral step lengths (see methods, Equation 1), which is a robust and clinically relevant measure conventionally used to characterize gait changes (31). During the practice session, participants experienced multiple bouts of alternating split (legs moving at different speeds) and tied belt (regular treadmill) walking (Figure 1A), which has been shown to accelerate locomotor savings in young adults (2, 32).

Figure 2A illustrates the time course of StepAsym during adaptation (split-belt) relative to baseline (tied-belt) in the pre- and post-practice sessions. Locomotor savings was observed by reduced gait perturbation (closer to zero StepAsym) in the post-practice (open circles) compared to the pre-practice session (filled circles, Figure 2A). We quantified locomotor savings with a savings index, defined as the change in StepAsym during early split-belt walking from pre-to post-practice as a proportion of the pre-practice StepAsym (see methods, Equation 2). We found a significantly smaller locomotor savings index in older adults than younger adults (two-sample t-test, t = -3.96, p < 0.001, Cohen’s d = -1.22; Figure 2B), suggesting that locomotor savings is reduced with age.

**Figure 2.**
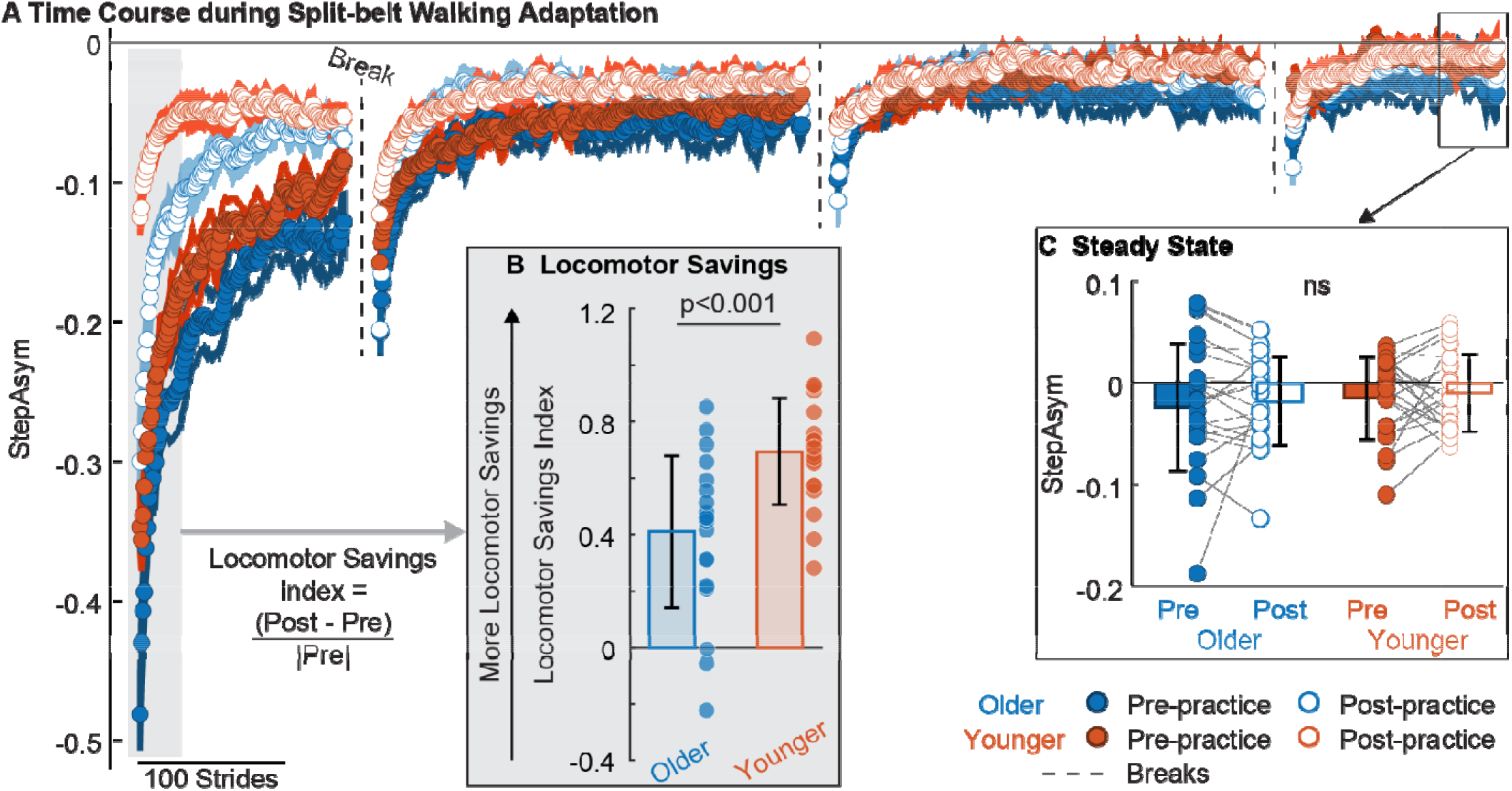
Step length asymmetry (StepAsym) time course (A) with inserts comparing locomotor savings (B) and steady state walking (C) between older and younger adults. (A) Group averaged time course of StepAsym during the adaptation condition (split-belt) in pre-(closed circles) and post-practice (open circles) sessions. All data is baseline (tied-belt) removed. Closer to zero values represent more symmetric (i.e., less perturbed) walking. Dot represents the average of 5 consecutive strides, and shaded area represents the standard error of each group. Both groups showed typical adaptation trajectories in both sessions (closer to zero StepAsym towards the end of adaptation), but the difference in early adaptation (grey shaded area) between pre- and post-practice sessions is larger in younger adults (red) compared to older adults (blue). (B) Locomotor savings. Bar plot represents the mean per group ± standard deviation. Dot represents the locomotor saving index value, calculated by equation shown in the figure. Older adults showed significantly less locomotor savings than younger adults. (C) StepAsym during steady state walking. Bar plot shows the mean per group ± standard deviation, and each dot corresponds to the average StepAsym during steady state walking (last 40 strides of adaptation) per participant. All participants reached the same steady state after adaptation in both sessions (as shown by the non-significant age and session effect).

Despite differences in locomotor savings, older and younger adults adapted to similar steady state walking behavior in both sessions (Figure 2C). StepAsym during the pre- and post-practice steady state periods (last 40 strides of adaptation) showed no significant main effect of age group (F_(1,40)_ = 0.22, p = 0.64), session (F_(1,40)_ = 0.13, p = 0.72), nor their interactions (F_(1,40)_ = 0.05, p = 0.82). These results suggest that older adults maintained the ability to adapt walking patterns with sustained exposures to a novel environment, but they saved less across days.

### Locomotor savings was correlated with higher attentional control of gait in older adults

To determine the role of attentional control of walking in locomotor savings, we evaluated the attentional control of gait using a well-established dual-task walking paradigm (20, 29). Participants walked at their comfortable pace around an oval while reciting every two (easy dual-task) or three letters of the alphabet (hard dual-task), with concurrent prefrontal cortical (PFC) activity recordings from a functional near-infrared spectroscopy (fNIRS) headband (Figure 1C). Similar results were observed in both tasks, and the following section will focus on the hard dual-task (see supplementary Fig. S1, Table S2, S3, for the easy dual-task). Attentional control of walking was quantified with an attentional gait index (33), which combines the PFC activation and motor and cognitive performance change from single-to dual-task walking (Equation 3 in methods). Higher values on the attentional gait index reflect greater reliance on attentional resources and reduced automatic control of gait (16, 18–21, 33). This indirect assessment was done as a first step to evaluate attentional control during walking due to technical challenges in acquiring reliable fNIRS-based PFC estimations during a dynamic adaptation task likes locomotor savings (see limitations).

We found that older adults have greater attentional control during walking than younger adults. This is demonstrated by the significantly higher attentional gait index in older than younger adults (z = 3.20, p = 0.001; Figure 3A). In the exploratory analysis, we found that this age difference in attentional gait index was driven by elevated PFC activation (z = 3.52, p < 0.001; supplementary Table S1), and a trending effect from greater motor performance cost (i.e., more reduction in motor performance from single- to dual-task; trending interaction, F_(1,40)_ = 3.13, p =0.08; supplementary Table S1) in older than younger adults.

**Figure 3.**
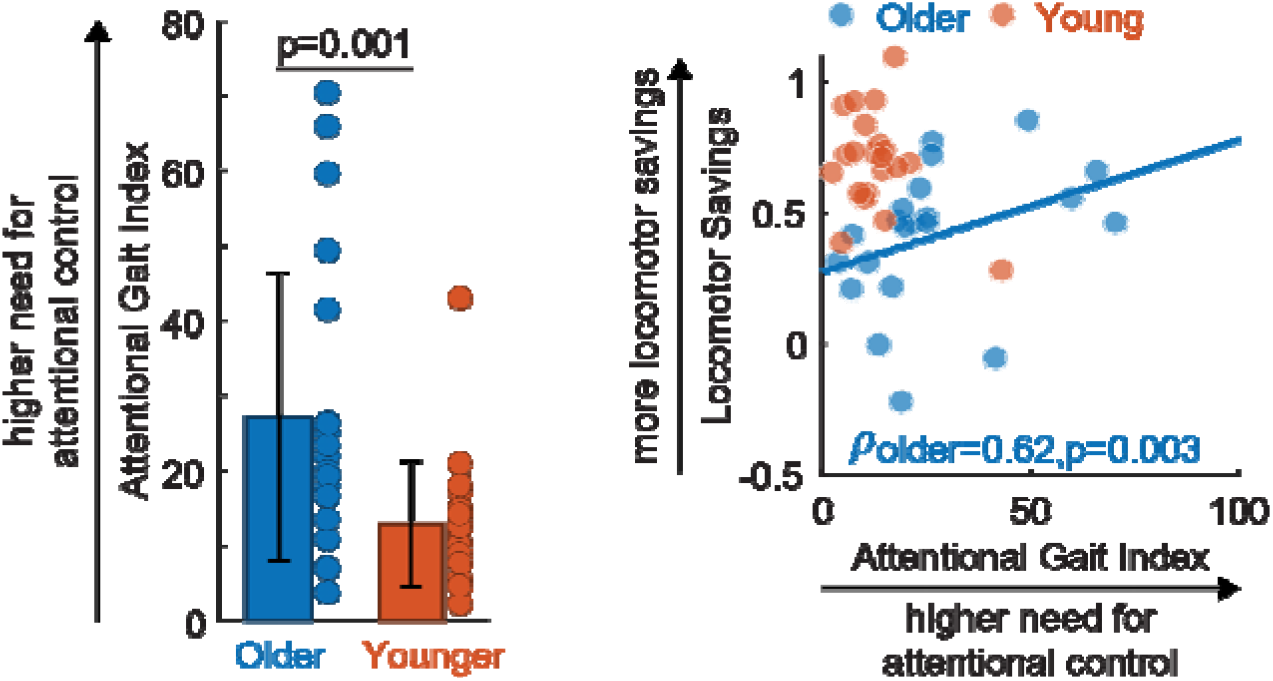
Attentional control of gait (measured by the Attentional Gait Index) comparisons between older and younger adults (A) and associations with locomotor savings (B). (A) Older adults showed significantly higher Attentional Gait Index values than younger adults. Bar plot represents the group mean per group ± standard deviation. Dot represents Attentional Gait Index (a metric combining prefrontal cortical activations and performance change from single-to dual-task walking) of each participant. (B) Scatter plots of Attentional Gait Index vs. locomotor savings. For visualization purpose, a linear regression line was displayed when we found a significant correlation. A significant positive correlation was observed in older adults.

More importantly, locomotor savings was positively correlated with attentional gait index in older (*ρ* = 0.62, *p* = 0.003; Figure 3B), but not in younger adults (*ρ* = -0.05, *p* = 0.82; Figure 3B). The association in older adults remained robust after controlling for age, sex, and self-selected overground walking speed (partial correlation *ρ* = 0.69, *p* = 0.001). To identify the factors driving the correlation between attentional gait index and locomotor savings in older adults, we performed exploratory analyses on the subcomponents of attentional gait index (PFC activity and performance), which are metrics that are conventionally reported in the literature (18–21, 34). We found that locomotor savings was correlated with motor performance cost (*ρ* = -0.45, *p* = 0.04), but not with cognitive performance cost (*ρ* = -0.20, *p* = 0.39) or PFC activity (*ρ* = 0.28, *p* = 0.22). Notably, the association between locomotor savings and motor performance cost was not influenced by single-task motor performance, as single-task motor performance was neither associated with locomotor savings (*ρ* = 0.04, *p* = 0.87) nor motor performance cost (*ρ*< 0.001, *p* = 1.00). The results indicate that attentional control of walking, as measured by attentional gait index, contributes to locomotor savings in older adults, and this relationship was mainly driven by motor performance cost.

Taken together, we found that older adults rely more on attentional control during walking than younger adults, and those with greater attentional control during walking exhibit greater locomotor savings. This relationship was not present in younger adults, suggesting an age-specific compensatory mechanism.

### Locomotor savings was weakly correlated with functional gait assessment

To explore if locomotor savings is relevant for functional mobility, we tested its relationship with the functional gait assessment (FGA), which evaluates an individual’s ability to perform multiple motor tasks while walking (35). We found a trending correlation where higher FGA score was associated with more locomotor savings (*ρ* = 0.38, *p* = 0.09, supplementary Figure S2). Of note, 18 out of 21 older adults in our sample scored above the fall-risk cutoff of 22/30 on the FGA (35– 37), suggesting we have a high functioning cohort (mean = 24.71±2.43, range = [18, 29]). Taken together, our results suggest that locomotor savings may only weakly reflect functional mobility in older adults without significant mobility or balance limitations.

### Locomotor savings in older adults was not correlated with cognitive abilities

Lastly, prior literature suggests cognitive processes are relevant for motor adaptation and savings in both reaching (6, 38) and walking (4). Therefore, we explored the relationship between locomotor savings and cognitive function using a comprehensive neuropsychological battery that assesses general mental status and major domains of cognitive functioning, including language, visuospatial ability, attention, memory, and executive functions (39–41). Among the domains tested, we observed only a trending relationship between the Repeatable Battery for the Neuropsychological Status (RBANS) (42) visuospatial index and locomotor savings where older adults with higher scores exhibited greater locomotor savings (*ρ* = 0.40, *p* = 0.07). Given the exploratory nature of these analyses and the sample size of our data these results should be interpreted with caution. Full neuropsychological test scores and correlation results are provided in supplementary Table S4.

## Discussion

We found that older adults can exhibit multi-day locomotor savings following practice, but to a less extent than younger adults. We also demonstrated that older adults have increased attentional control of walking, and higher attentional control of walking is linked to greater locomotor savings in older adults. We argue that with aging, different aspects of motor behaviors (such as locomotor savings and dual-task walking) become linked due to increased reliance on attentional compensations following age-related decline in the shared neural substrates. Future work is needed to test how this relationship changes with different cognitive abilities such as in older adults with cognitive impairment.

### Older adults can save and recall learned locomotor patterns with practice

In this study, we demonstrated that older adults exhibited locomotor savings to a lesser extent than younger adults. Mixed findings have been reported about the influence of aging on savings in reaching (25, 43, 44) and walking (13, 14, 45–47) where older adults show either similar or reduced savings depends on the task, protocol, and assessment intervals.

The main factor that could explain these discrepancies is our use of a structured practice session, which is known to favor locomotor savings (2, 4, 32, 48). We used a practice protocol that included abbreviated bouts of novel walking interleaved with regular walking on the practice day (Figure 1A). These transitions provided repeated experiences of movement errors, a critical driver for savings in younger adults (5, 48, 49). We extend previous research by demonstrating that multiple movement errors also effectively induce locomotor savings in older adults. Unlike prior multi-day studies (12, 14, 25) our structured practice likely enhanced memory consolidation, allowing older adults to show locomotor savings.

The second factor distinguishing our study from previous is the controlled passage of time between learning and re-learning (13, 50, 51). Previous work has reported that older adults forget a newly learned walking pattern more and showed less retention than younger adults (13, 51, 52), suggesting faster decays of recent motor memories in older than younger adults. In models comprising fast and slow learning components, savings is typically attributed to the slow component (3, 53, 54), and older adults often show reduced retention of the slow component (55). Taken together, while older adults may demonstrate savings within a single session where memory decay is incomplete (43–46), over longer intervals, such as the six-week in the previous work, older adults’ memory of the locomotor patterns may decay more until no savings remain (12, 13). In our study, intervals between sessions were carefully controlled, with an average interval of 3.98 days between initial exposure (pre-practice session) and the locomotor savings assessment (post-practice session). The controlled time interval and the inclusion of a practice session may have allowed us to detect the age-related difference in locomotor savings across days.

In sum, we demonstrate that both older and younger adults can exhibit multi-day locomotor savings after practice and controlled short time interval. Nonetheless, even under these controlled conditions, older adults showed less locomotor savings than younger adults, suggesting age-related limitations in long-term motor memory savings.

### Older adults retain the flexibility to adapt in a novel environment

Despite differences in locomotor savings, older adults reached the same steady state behavior as younger adults at the end of the adaptation. This indicates that although older adults were more perturbed during early adaptation, they were able to adapt and achieve similar gait symmetry as younger adults after a sustained exposure to the split-belt treadmill environment. This is consistent with prior studies in walking (45, 46, 56–58) and reaching (43).

The hierarchical computational model of locomotor adaptation and savings (59) provides a framework for interpreting these findings. The model comprises three key components: (1) a reinforcement learner that improves performance, (2) a feedback controller that stabilizes the body, and (3) a memory module that stores previously learned walking strategies. The age-related decline in locomotor savings may emerge from deficits in one or more of these components.

Dopaminergic reinforcement learning system is relevant for savings in reaching (48, 60, 61) and dopaminergic neurotransmission has been associated with walking adaptation in aging (62, 63). Although older adults eventually achieved similar asymmetry to younger adults, they took longer to counteract the initial perturbation (13, 46, 56, 58, 64). Deficits in the dopaminergic reinforcement learner can contribute to the slower learning within the session and reduced savings across sessions.

Alternatively, a weakened feedback controller could contribute to poor movement adjustments when encountering a novel environment, resulting in more perturbed movement during early split-walking in older adults across all sessions (Figure 2A, blue circles for older adults were lower than red circles for younger adults in both sessions). This aligns with balance (65–67) and walking (64) literature showing reduced reactive feedback correction in older adults. A less efficient feedback controller would result in larger errors during both learning and relearning (savings).

Lastly, our results could indicate a faulty memory module in older adults that limit the update and retrieval of the corresponding motor memory when re-encountering the split environment (59), resulting in poor locomotor savings (Figure 2B). This is consistent with the fast and slow learning components model, which attributed savings to a biased slow learning component during re-learning (3, 53, 54). Older adults exhibited reduced retention in this slow component (51, 55), which could result in a smaller bias during re-learning and therefore diminished locomotor savings observed in this study.

Note that these hypotheses are not mutually exclusive. Future computational work is needed to distinguish the specific components underlying the age-related differences in locomotor savings.

### Attentional compensations can be a strategy for locomotor savings

We observed that higher attentional control of walking is correlated with greater locomotor savings in older adults. This suggests that cognitive compensation, typically indicative of reduced automaticity (16, 18–21), can be advantageous for complex walking behaviors like locomotor savings (4, 68). This relationship may stem from shared neural correlates such as the subcortical-frontal (e.g., basal ganglia and cerebellum to primary motor cortex) networks (45, 69, 70). These structures are susceptible to age-related changes in white matter hyperintensity volume and functional connectivity (71–73). Consequently, older adults may increasingly recruit PFC-mediated attentional and executive resources as a compensatory mechanism for challenging walking (16). Supporting this view, executive cortical-striatal network resting-state connectivity has been linked to mild Parkinsonian signs (72, 74). In healthy aging, older adults with higher attentional control of gait in dual-task walking may be more inclined to use cognitive strategies in other challenging motor tasks such as locomotor adaptation, and this strategy may facilitate locomotor savings.

We were unable to measure PFC activity concurrently with locomotor savings due to technical limitations and we inferred the role of attentional control in locomotor savings using attentional control of walking measured by behavior and PFC activation in dual-task walking. Future multi-modal imaging studies are necessary to provide direct evidence for the neural correlates of locomotor savings. Furthermore, the relationship between locomotor savings and attentional control of walking was most robustly observed in the attentional gait index, highlighting the importance to integrate performance when interpreting fNIRS-based PFC activities, as shown in our previous work (33).

### Attentional gait index revealed reduced automaticity and inefficient use of attentional resources during challenging walking in older adults

We found that older adults exhibited increased attentional control of walking, as indicated by higher attentional gait index and higher PFC activation compared to younger adults across tasks. These findings align with prior work showing reduced gait automaticity in aging (16, 21) where older adults showed elevated PFC activation or greater dual-task performance cost (16, 18–21). Notably, our older adult cohort recruited more PFC yet performed worse than younger adults, suggesting the extra PFC recruitment was inefficient. Taken together, our results indicate that older adults had reduced gait automaticity, increased reliance on attentional resources, but limited benefit from this compensatory recruitment during dual-task walking.

### Other metrics of functional mobility are needed to determine the relationship between mobility and locomotor savings

Our exploratory analysis revealed only a weak correlation between locomotor savings and dynamic balance measured by the Functional Gait Assessment (FGA). However, this does not mean locomotor savings is irrelevant for mobility. The lack of association may be due to a ceiling effect in our cohort’s FGA scores. Prior research has suggested that higher levels of self-reported recent physical activity was related to better locomotor adaptations (57), and gait adaptability is similar between physical activity matched older and younger adults (75). This warrants future research to use more comprehensive mobility metrics, such as accelerometry or GPS-based objective measures of real-world physical activities and community mobility (76) and examine their associations with locomotor savings.

### Cognitive assessment results were not directly associated with locomotor savings

Most neuropsychological test results in our battery were not correlated with locomotor savings. While cognitive processes are relevant for savings in younger adults, such as explicit re-aiming (6), explicit recall of perturbation size (4), and visual field dependence (77), evidence in older populations remains sparse. One study showed no relation between adaptation rate and a variety of neuropsychological test results in older adults (78), while Sasikumar and collegues (79) reported that better overall cognitive scores were related to adaptation and aftereffects in patients with Parkinson’s disease. Taken together, these findings suggest that neuropsychological test scores may only predict locomotor adaptation and savings in more impaired populations (e.g., Parkinson’s disease patients), but not in older adults with little mobility or cognitive limitations like in the current sample. Alternatively, the relationship between cognitive processes and locomotor savings may only emerge under conditions that pose sufficient challenge to the nervous system in both motor and cognitive domains simultaneously, such as in dual-task walking, rather than through static neuropsychological assessments.

### Limitations

The main limitation of this study is that attentional compensation was not measured concurrently with locomotor savings. Reliable fNIRS-based PFC estimations require multiple repetitions of a task to ensure a sufficient signal-to-noise ratio. However, since repeated exposures to split-belt perturbations inherently alter adaptation behavior, each encounter represents a dynamic, non-stationary state rather than a simple repetition. To address this challenge, future studies could develop new fNIRS analysis supporting continuous measurement of a dynamic condition, use indirect measures of attention (e.g., arousal), or incorporate imaging modalities with better temporal resolution (80), such as electroencephalogram (EEG).

### Clinical Implications

The relationship we identified between locomotor savings and attentional control of walking in older adults is of clinical relevance because it reflects cognitive compensation strategies in walking, which is a strong indicator of dementia risks (81). It is important to investigate how this relationship changes when cognitive compensation is compromised due to neurodegeneration, such as in older adults with cognitive impairment. Understanding these mechanisms will also help identify older adults who are less likely to retain clinical gait interventions due to underlying cognitive decline.

## Conclusions

In conclusion, we found that practice effectively induced multi-day locomotor saving in older adults, though to a lesser extent than in younger adults. We also found a moderate relationship between attentional control of walking and locomotor savings in older adults, but not in younger adults. This suggests that locomotor savings relies more on higher-order processes such as attention in older ages, and the attentional compensation might support older adults to save and recall from prior experiences. Future work is needed to directly examine the neural substrates underlying this relationship and determine if the relationship persists when compensatory neural resources are diminished, such as in older adults with cognitive impairment.

## Materials and Methods

### Participants

Older (>= 65 years old) and younger (19-40 years old) adult participants were recruited through flyers posted in the community and through research registries including the Pitt+Me Research Recruitment Program and the Pittsburgh Pepper Community Research Connection. All participants could walk unassisted, did not take medications that change cognitive functions, and had no major neurological conditions, severe visual impairment, or unmanaged orthopedic conditions that would interfere with walking. Older adults were further screened for signs of cognitive impairments as indicated by a MoCA score less than 24. A total of 25 older adult participants were recruited; three did not meet the MoCA inclusion criteria and one dropped out, resulting in 21 older adult participants for all data analysis. Sex matched younger adults were recruited with one drop out, resulting in 21 participants for data analysis. Participant demographics are included in Table 1. The Institutional Review Board at the University of Pittsburgh approved the study. All participants gave written informed consent.

### General Experimental Protocol

All participants performed dual-task walking while wearing a functional near-infrared spectroscopy (fNIRS) headband to evaluate attentional control of walking (Fig. 1C) before exposure to the split-belt treadmill. Then, participants completed three sessions of split-belt treadmill walking on separate days, pre-practice, practice, and post-practice (Fig. 1A). Each session was one or two days apart (between pre-practice and practice sessions: 1.67 ± 2.34 days (mean±standard deviation), between practice and post-practice: 1.31 ± 0.56 days). The evaluation for attentional control of walking was completed on the same day before the first split-belt exposure for all younger adults, resulting in three total experimental sessions in younger adults. Older adults visited the lab one more time one-or two-days prior to the first split-belt treadmill walking session, during which they completed a comprehensive neuropsychological exam, attentional control of walking assessment, and Functional Gait Assessment (FGA). This extra visit was added to avoid overburdening the participants at any single session. Importantly, in both groups, attentional control of walking was always evaluated before any treadmill walking to characterize participants’ baseline abilities.

### Protocol to Evaluate Locomotor Savings

Locomotor savings of a newly learned walking pattern was evaluated across multiple exposures to a split-belt treadmill where the two legs moved at different speeds. The protocol consists of three sessions (Fig. 1A): pre-practice, practice and post-practice. All sessions consist of baseline, adaptation, and washout conditions. Detailed treadmill speed profile for the whole protocol can be found in supplementary material Figure S3.

Baseline was collected during both overground and treadmill walking. During overground baseline, participants walked back and forth on a 9.2m walkway at a self-selected speed for 150 strides excluding turns. A stride was defined as two consecutive heel-strikes of the same leg and heel strikes were identified in the kinematic data as the times of the ankle marker’s greatest forward excursion (13, 82, 83). During treadmill baseline, participants walked on an instrumented treadmill (Bertec, Columbus, OH, United States) with both legs moving at the same speed (tied-belt) at 0.5m/s, 1.0m/s, and 0.75 m/s for 150 strides each. Then, during adaptation, participants walked on the treadmill with the dominant leg moving at 1m/s and non-dominant leg moving at 0.5m/s (split-belt). Participants experienced this novel situation in a single block of 900 strides in split-belt in “pre” and “post-practice” sessions (Fig. 1A). In contrast, during the “practice” session, participants experienced five bouts of 250 strides with the treadmill belts split interleaved with four bouts of 50 strides with the treadmill belts tied at 0.75m/s (Fig. 1A). Next, during washout, participants walked 450 strides at their self-selected speed overground and then 150 strides with tied-belt treadmill at 0.75m/s. The overground and tied-belt walking were included to wash out the adaptation effect from the session.

After treadmill washout condition, participants experienced two short perturbations in opposite directions (supplementary material Figure S3). Overground washout and short perturbations were performed for a different aim in the larger study to evaluate adaptation in muscle activities (84, 85) and were not used in the analyses presented in this paper. We included the information here to promote reproducibility. However, we believe these protocol design choices do not impact our overall conclusions.

Participants took sit-down resting breaks during regular intervals to limit fatigue (dashed lines in Fig. 1C). The breaks were administered at the same location with the same durations for all participants (break duration for older adults: 5.35±0.58 mins (mean±standard deviation), younger adults: 5.16±0.41 mins). Participants were instructed to not move their feet during the breaks.

### Locomotor Savings Data Collection

Kinematic data during split-belt walking was collected with Vicon Motion Systems (Oxford, United Kingdom) at 100Hz. We placed passive reflective markers bilaterally over the hip (greater trochanter) and ankle (lateral malleolus). Gaps in the raw kinematic data due to marker occlusions were filled first with a quintic spline interpolation (Woltring; Vicon Nexus Software, Oxford, United Kingdom) and remaining gaps were filled manually using Vicon’s spline, pattern, and cyclic fill tools when appropriate as done in previous studies (12).

### Locomotor Savings Outcome Measures

Behaviors on the treadmill were quantified by step length asymmetry (StepAsym, Figure 1B), defined as the normalized difference in bilateral step lengths (Equation 1). The normalization allows fair comparison among individuals with different stride length. A zero value for StepAsym indicated that both steps were the same length, i.e., symmetric walking. Step length is quantified by the distance between the ankle markers at heel strike, specifically fast step length was measured during fast leg heel strike and vice versa for the slow leg.

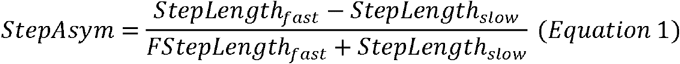

Locomotor savings was quantified by a locomotor savings index, defined as the difference in StepAsym during early adaptation (first 30 strides in adaptation) between the pre- and post-practice sessions, normalized by pre-practice (Equation 2). This normalization allowed us to quantify locomotor savings relative to individual’s baseline ability to counteract the split-belt perturbations. The average of the first 30 strides was chosen as a model-free approximation for the adaptation rate without assuming any specific patterns of the stride-dependent relationship (2, 26). Higher values on the locomotor savings index indicate less perturbed movements when re-exposed to split-belt walking during the post-practice compared to pre-practice session, which means more locomotor savings.

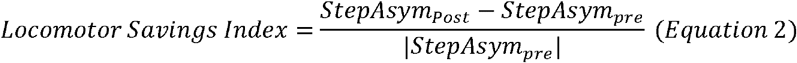

StepAsym during pre-practice was always negative for all participants. So, absolute value was taken on the denominator such that the sign of the locomotor savings index is determined by the differences between post and pre-practice StepAsym and a positive locomotor savings index represents improved StepAsym (closer to zero gait) in post-relative to pre-practice.

### Protocol to Evaluate Attentional Control of Walking

Attentional control of walking was evaluated using a well-established dual-task walking paradigm (20, 29). Participants walked at their comfortable pace along on oval path with a ∼5.85-meter straight walkway on each side, while reciting every two (e.g., A, C, E, etc.) or three letters (e.g., A, D, G, etc.) of the alphabet and wearing a functional near-infrared spectroscopy (fNIRS) headband to measure prefrontal cortical (PFC) activity (Fig. 1C). Reciting every two or three letters while walking was considered the easy or hard dual-task conditions, respectively. Two levels of difficulties were included to allow calculation of an attentional gait index as shown in our previous work (33). Participants also performed a motor single-task, walking without any secondary task, and two cognitive single-tasks, standing while reciting every two (easy) or three (hard) letters of the alphabet (Fig. 1C). The single-task conditions allowed the quantification of performance change from single-to dual-task walking.

We used a block design with six trials. Each trial includes all three single-tasks and two dual-tasks in a pseudo random order. The task order within a trial was kept the same across all participants, but the order of the trials was randomized. Each task was performed for 20 seconds, and a standing rest preceded every task. During the standing rest, participants were instructed to stand while counting silently from one to allow the hemodynamic response to return to rest level while limiting distraction or practice of the cognitive task during rest (86).

Instructions for each condition were given by a computerized voice using a customized MATLAB script that automatically controls task timing. Before starting the session, we confirmed that all participants could hear the instructions clearly. Participants were instructed to walk at their comfortable pace while doing the cognitive task with no specific instruction about which task to prioritize. A practice trial was performed up to three times to ensure the participant understood the task instructions (repeated 1.02±0.15 (mean±standard deviation) times).

### Attentional Control of Walking Data Collection and Data Processing

Motor performance was quantified by gait speed (speed, m/s) using motion data from Vicon Motion Systems (Oxford, United Kingdom). Motor performance during dual-task was calculated as the total Euclidian distance in the x-y (transverse) plane traveled by the average position of the hip markers (greater trochanter) divided by the time the marker was moving. The hip markers’ motion data was collected using the same system and processed with the same pipeline as the locomotor savings data (see *Locomotor Savings Data Collection* section above).

Cognitive performance was measured by rate of correct letters of alphabet generated (rate, letter /s). An experimenter transcribed the alphabet generated by the participant during the experimental session and the rate of correct letters of alphabet generated was calculated by a customized MATLAB script during post-processing.

PFC activity was measured from an eight-channel continuous wave functional near-infrared spectroscopy (fNIRS) headband (Octamon, Artinis Medical Systems, Netherlands). The fNIRS headband consisted of two detectors and eight sources with a source and detector pair distance of 35mm. The fNIRS headband covered both the left and right PFC regions, specifically Brodmann areas 9, 44, 45, and 46 (34). The center of the fNIRS headband was aligned with the center of the participant’s nose, and the bottom of the headband was just above the eyebrow (87). The headband was not removed between trials to maintain stable placement.

fNIRS quantifies changes in the blood oxygenation based on distinct light absorption properties of the oxygenated (Hbo) and deoxygenated (Hbr) blood, which provides a proxy for cortical activities. Near-infrared light transmitted at 850□nm and 760□nm was used to detect changes in Hbo and Hbr. Data were sampled at 10□Hz and collected by the OxySoft software (Artinis Medical Systems, Netherlands). No short separation channel was available in the equipment, and physiological and extracortical noises were addressed statistically (see below) using the NIRS Brain AnalyzeIR toolbox (88) in MATLAB 2021a (Mathworks, Natick, Massachusetts). Hbo typically has a stronger signal-to-noise ratio (69) and therefore was used for the analysis presented in the main paper. The same analysis can be performed with Hbr (supplementary Table S5, S6).

fNIRS data was processed with the NIRS Brain AnalyzeIR toolbox (88) following the procedures described in our previous work (33). Data for a given task were excluded from processing and analysis (0.01% of the tasks) if there were experimental errors or if the participant clearly violated the protocol (e.g., walking during a standing task). The fNIRS recording for each trial was first trimmed to keep only 2□s of the data before and after the first and last task of a trial to reduce global baseline noise. Channels with a variance less than 10^-9^ in a 5-s moving window, likely due to equipment malfunction, were identified as flat (0.01% of the data) and removed from analysis. Light intensity was converted to optical density and then converted to Hbo measurements using the modified Beer–Lambert law with a partial path length factor of 0.1 (34). The time series data for each source–detector pair was used to fit a general linear model (GLM). The design matrix of the GLM was the convolution of stimulus timing, duration, and a canonical hemodynamic response function (34). To minimize motion and physiological artifacts, the model was solved with an autoregressive pre-whitening iteratively reweighted least square approach (88). In brief, the autoregressive filter is a statistical method to alleviate physiological noise and motion artifacts. The iterative reweighted least square approach further downweighs large motion artifacts (87– 89). A Student’s t-test was then performed on the regression coefficients, and the t-score represents the changes in Hbo in each task compared to the standing rest while counting silently immediately before. Higher values represent more neural activity in the experimental task relative to standing rest. Notice that this analysis approach also alleviates the need to adjust for age-dependent differential pathlength factor because the mean and variance from the GLM will scale accordingly. The results across six trials and eight channels covering the whole PFC were combined using a weighted average that accounts for the covariance using the toolbox.

### Attentional Control of Walking Outcome Measure

Attentional control of walking was measured by attentional gait index that combines PFC activations and dual-task performance (Equation 3, ref Liu 2024).

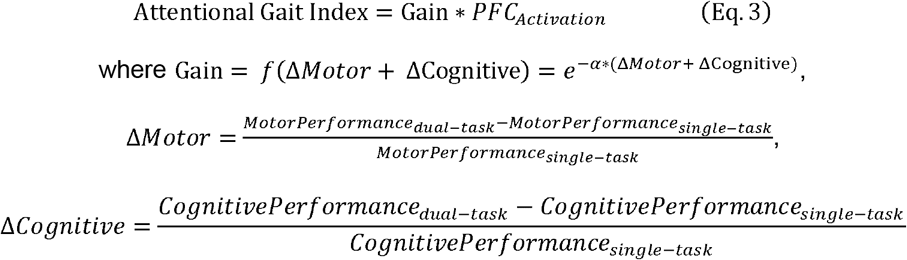

The optimization of the hyperparameter α and index calculation followed the same procedure as described in our previous work (33). The definition of ΔMotor and ΔCognitive is similar to dual-task cost in existing literature (90, 91) and the normalization relative to single-task allowed fair comparisons among individuals with different baseline motor and cognitive performances.

The attentional gait index is a monotonic axis that quantifies the need for attentional control during challenging walking, which has been shown to be more informative than PFC or dual-task performance alone (33). Higher attentional gait index values represent greater PFC recruitment and poorer performance in dual-task relative to single-task walking, which are indicative of greater reliance on attentional resources and reduced automatic control of gait (16, 18–21, 33).

### Self-selected overground walking speed

Self-selected overground walking speed was calculated as the average speed walking back and forth on a straight 9.2m walkway excluding turns for 150 strides during the baseline overground shown in Figure 1A.

### Functional Gait Assessment Protocol

The Functional Gait Assessment (FGA) was scored in older adults on an ordinal scale for 10 items with a max score of 30 and higher score indicates better balance function (30). The 10 items are: gait on level surface, change in gait speed, gait with horizontal and vertical head turns, gait with 180° pivot turn, stepping over obstacles, gait with narrow base of support, gait with eyes closed, backwards gait and stairs (30). Scoring of each of item is done on a 4-point ordinal scale ranging from 0 indicating severe impairment to 3 indicating no impairment. The assessment was always administered by the same experimenter (SL) to reduce inter rater bias.

### Neuropsychological Assessment Battery

Older adults completed a comprehensive neuropsychological assessment in their first visit to the lab. The battery was designed and reviewed by a clinical neuropsychologist (AW) and the tests were administered and scored by trained psychometrists. General cognitive status was measured with the Montral Cognitive Assessment (MoCA). Verbal ability/literacy was measured with the Wide Range Achievement Test Fifth Edition (WRAT5) (92). Overall cognition was measured with the Repeatable Battery for the Assessment of Neuropsychological Status (RBANS) (42), which includes tests that factor onto five cognitive domain index scores: Attention (Digit Span, Coding), Language (Picture Naming, Semantic Fluency),Visuospatial (Figure Copy, Line Orientation), Immediate Memory (List Learning, Story Learning), and Delayed Memory (List Recall, Story Recall, Figure Recall). A supplemental language test, the action verbal fluency test (93), was administered because the task is sensitive to basal ganglia function and may be relevant in locomotor savings (94). Executive function was measured with subtests of the Delis-Kaplan Executive Functioning System: Color Word Interference, Verbal Fluency, and Trail Making Test. Since these subtests were co-developed and co-normed, we combined them into a composite executive function score by averaging scaled scores from Color-Word Interference condition 3, Color-Word test 4, Trail Making Test condition 4, and Verbal Fluency conditions 1, 2, and 3 total correct. Similarly, cognitive switching was quantified by a composite score averaging Color-Word Interference test condition 3, Trail Making Test condition 4, and Verbal Fluency condition 3 category switching scaled scores from D-KEFS (95).

### Power Analysis

This study aims to evaluate the correlation between attentional control of walking and locomotor savings in older adults. A sample of 21 older adults achieves 80% power to detect a moderate (96) correlation (r = 0.57) at α = 0.05. To compare across age, a sex-matched sample of 21 younger adults was also recruited. This total sample size provides 80% power to detect a large (97) between-group difference in locomotor savings (Cohen’s d = 0.89) at α = 0.05. All power analyses were conducted in R (version 4.5.1) using the pwr package.

### Statistical analysis

In the primary analysis, t-test with equal variance was used to compare locomotor savings index between older and younger adults after checking that normality (Shapiro-Wilk test, p = 0.24 and p = 0.58 for older and younger adults respectively) and homoscedasticity were satisfied (Levene’s absolute test, p = 0.18). A Spearman’s correlation was performed for each age group to quantify the relationship between locomotor savings and attentional control of walking (measured by the attentional gait index), because we were interested in quantifying the monotonic relationship and the attentional gait index was not always normal.

In the exploratory analysis, due to violations of normality, non-parametric tests were performed. To compare age-related differences in attentional gait index and PFC activation, Wilcoxon’s rank sum test was performed. To compare difference in motor or cognitive performance in single- and dual-task walking as well as differences in steady state split-belt treadmill walking, aligned rank tests were performed (98–100) with age as the between-subjects factor and task (single- or dual-task for motor and cognitive performance) or session (for steady state walking) as the within-subjects factor. When a significant correlation was found in the primary analysis, we performed exploratory Spearman’s correlations to identify if the relationship was driven by any subcomponents of the attentional gait index. Spearman’s correlation was also performed to explore the relationship between locomotor savings, functional gait assessment, and neuropsychological test results.

A significance level of alpha = 0.05 was used for all analysis. Aligned rank test was performed in R (version 4.5.1) and all other tests was performed in MATLAB 2021a (Mathworks, Natick, Massachusetts).

## Acknowledgments

This study was supported by National Institutes of Health 1R01AG089175-01, National Science foundation Career Award 2419849, and Pittsburgh Pepper Center P30AG024827. SL is supported by the National Institutes of Health R90DA060340-02S1. AW is supported by the National Institutes of Health (K23AG076663). EB is supported by the National Institutes of Health (T32AG021885).

The authors thank Devin Stafford, Elan Albalak, Emma Stein, Shaoyi Li, and Shaina Patel for their tremendous help during data collection. The authors thank Dr. Dulce M. Mariscal, Dr. Krista Fjeld, Marcela Gonzalez-Rubio, and Jiwon Choi for their insightful feedback on the manuscripts and help during data collection. The authors thank Nathan Brantly and Adwoa A. Awuah for their feedback on the manuscripts. The authors thank Dr. Ted Huppert for his feedback on the fNIRS statistical analysis.

