## SupplementaryMaterials for "Locomotor savings relies on attentional control of walking in older, but not younger adults"

Figures

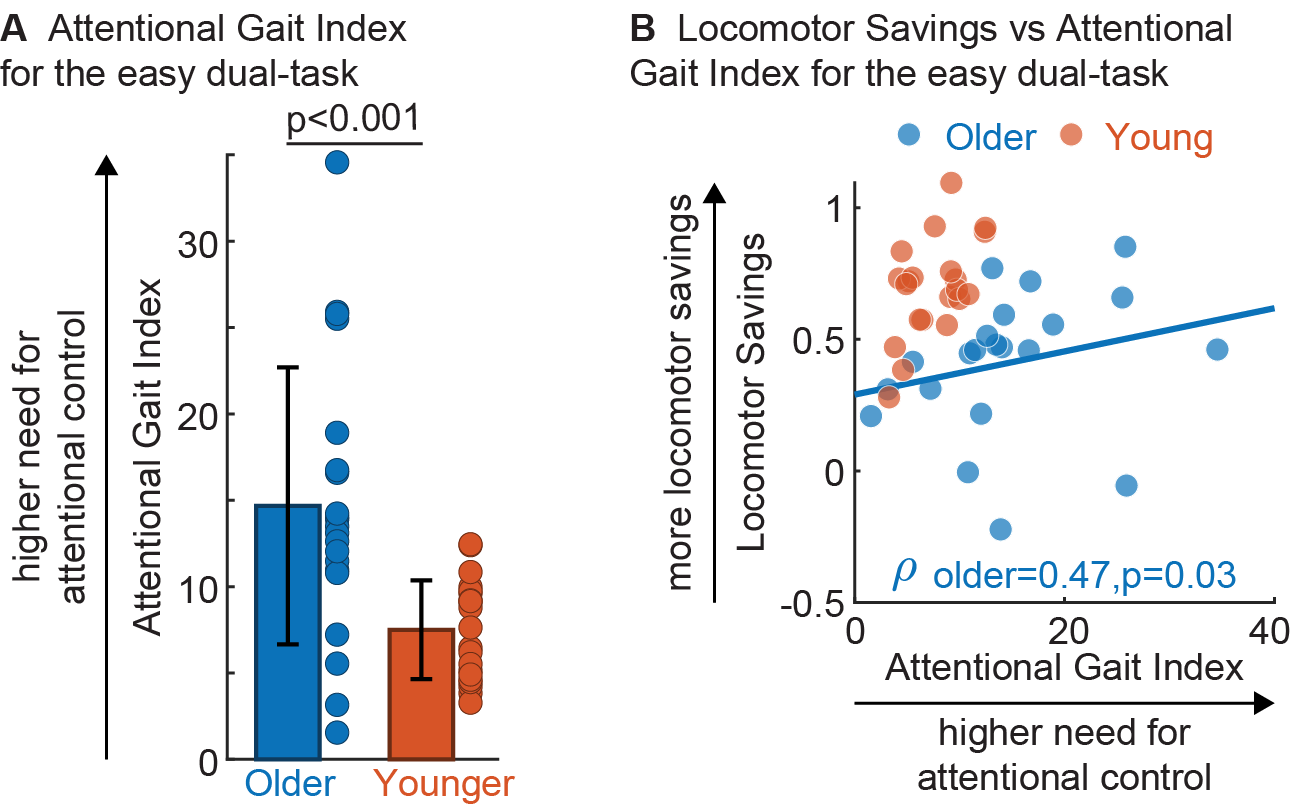

Fig. S1. Attentional control of gait comparisons between older and younger adults (A) in the easy dual-task condition (walk while reciting every two letters of the alphabet) and associations with locomotor savings (B). (A) Consistent with the findings in the hard single task in the main paper (Fig. 3), older adults showed significantly higher attentional gait index values than younger adults. Bar plot represents the group mean per group ± standard deviation. Dot represents attentional gait index of each participant. (B) Scatter plots of attentional gait index (x-axis) and locomotor savings (y-axis). For visualization purpose, a linear regression line was displayed when we found a significant correlation. Consistent with the findings in the easy single task presented in the main text, a significant positive correlation was observed in older adults only.

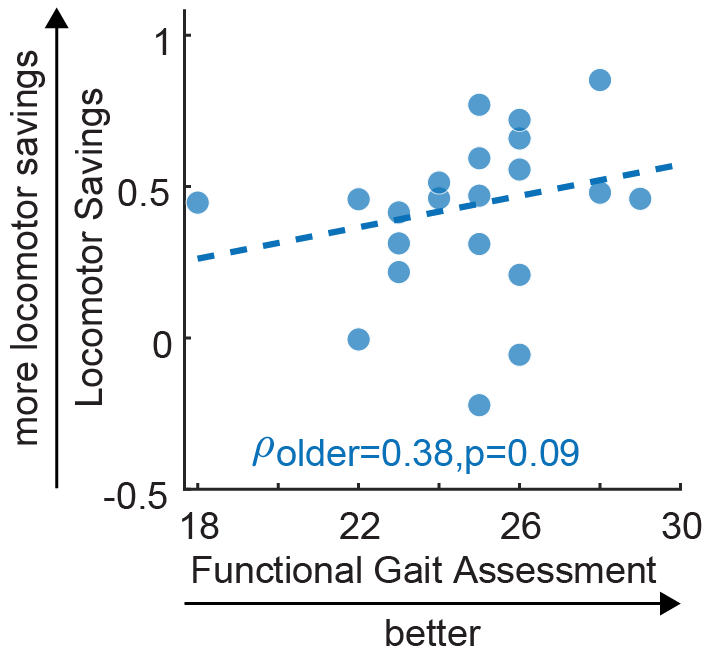

Fig. S2. Scatter plot of functional gait assessment (FGA) scores vs. locomotor savings in older adults. For visualization purpose, a dashed linear regression line was displayed when we found a trending correlation. A trending correlation was observed where older adults with higher FGA score showed more locomotor savings.

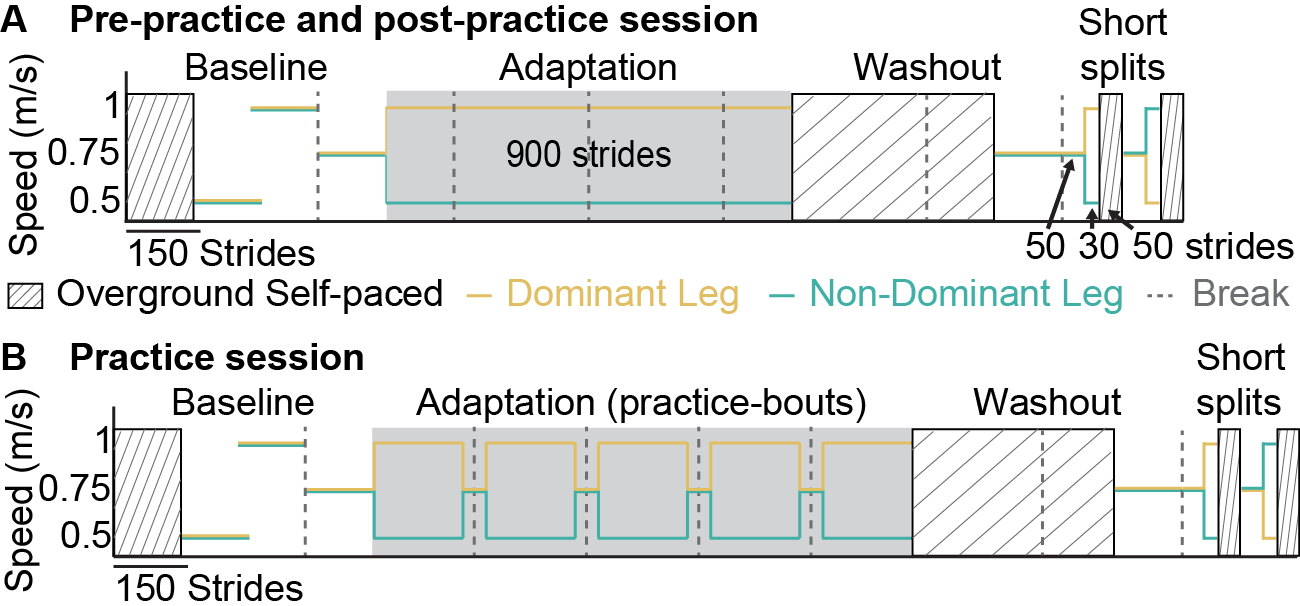

Fig. S3. Treadmill speed profile for the entire protocol. Green and yellow lines represent belt speed of the dominant and non-dominant leg respectively. Sit-down breaks were taken every 300 strides (dashed line). All participants experienced three sessions of split-belt treadmill walking as described in the methods of the main text (Figure 1A). Each session started with a baseline condition including 150 strides of overground walking at self-selected speeds (strip filled), 150 strides of slow, fast, and medium speed tied-belt walking. Then without a break, the belt under the dominant leg (green line) started to move at 1m/s while the belt under the non-dominant leg (orange line) slowed down to 0.5m/s abruptly. Participants walked in this split-belt condition for 900 strides in pre- and post-practice session or for 250 strides interleaved with tied walking of 50 strides in the practice session. Immediately after adaptation, they walked at self-selected speed overground for 450 strides for washout. Then participants were washed out again on the tied treadmill at 0.75m/s for 150 strides. Next, participants experienced a short-splits condition, which includes two perturbations in opposite directions. Specifically, participants walked with both belts moving at 0.75m/s for 50 strides, then they walked for 30 strides with the belts split in the same direction as the adaptation (i.e., the dominant and non-dominant leg walked at 1m/s and at 0.5m/s respectively), and immediately after they walked overground at their self-selected speed for 50 strides. This series was then repeated with the belt split in the opposite direction. Specifically, participants walked 50 strides at 0.75m/s on the tied-belt treadmill, then they walked for 30 strides with the treadmill belts split in the opposite direction as the adaptation, where the dominant leg now moved at 0.5m/s and non-dominant leg moved at 1.0m/s. Finally, they walked 50 strides at their self-selected speed overground again. The short-splits condition is collected to evaluate muscle activity adaptation (1, 2) as described in previous studies for a different aim of the larger study and not used in the analysis presented in this paper. We believe these epochs do not impact our conclusion presented in the main text. Importantly, both age groups experienced the exact same protocol.

Tables

Table S1. Raw values and between age-group comparisons of the attentional gait index (AGI)’s subcomponents in the hard dual-task condition described in the main text. PFC activation was compared with the Wilcoxon rank sum test. Motor and cognitive performance were compared with aligned rank tests, non-parametric method for two-way ANOVA.

| **PFC Activation** | | | | | | |
| --- | --- | --- | --- | --- | --- | --- |
|  | | **Older Adults**  **Mean (SD)** | **Young Adults**  **Mean (SD)** | **Test stats** | | |
| PFC activation during hard dual-task | | 4.21 ± 4.13 | -0.29 ± 2.34 | z = 3.52, p < 0.001 | | |
| **Performance** | | | | | | |
| **Domain** | **Task** | **Older Adults**  **Mean (SD)** | **Young Adults**  **Mean (SD)** | **Age Group Effect** | **Task Effect** | **Group x Task Interaction** |
| Motor  (speed, m/s) | Walk | 0.97 ± 0.13 | 1.01 ± 0.13 | F (1,40) = 1.55, p = 0.22 | F (1,40) = 91.59, p < 0.001 | F (1,40) = 3.13, p = 0.08 |
|  | Hard dual-task | 0.82 ± 0.14 | 0.91 ± 0.15 |  |  |  |
| Cognitive (rate, letters/s) | Standing cognitive task (Easy) | 0.50 ± 0.12 | 0.58 ± 0.10 | F (1,40) = 5.53, p = 0.02 | F (1, 40) = 85.80, p < 0.001 | F (1, 40) = 0. 42, p =0.52 |
|  | Hard dual-task | 0.41 ± 0.10 | 0.49 ± 0.14 |  |  |  |

Table S2. Subcomponents of attentional gait index (AGI) in the easy dual-task that drives the correlation between AGI and locomotor savings in older adults as observed in Figure S1B. Significant Spearman’s correlation was highlighted in red. Consistent with the findings in the easy dual-task shown in the main text, the relationship between AGI and locomotor savings in older adults was driven by motor performance change.

| AGI subcomponents in easy dual-task | Older Adults |
| --- | --- |
| PFC activation (Hbo) | $\rho=0.23, p=0.$31 |
| $\Delta$Motor | $\rho=-0.49, p=0.$03 |
| $\Delta$Cognitive | $\rho=-0.22, p=0.34$ |

We confirmed that the relationship in the easy dual-task remained robust after partial controlling for age, sex, and self-selected overground walking speed (partial Spearman’s correlation $\rho=0.49, p=0.04$). Furthermore, the association between locomotor savings and motor performance cost was not influenced by single-task motor performance, as single-task motor performance was neither associated with locomotor savings$(\rho=0.04, p=0.87$, as shown in the main text) nor motor performance cost ($\rho=0.04, p=0.85$, i.e., faster single task walkers did not always slow down more in dual-task).

Table S3. Raw values and comparisons of attentional gait index’s subcomponents in the hard dual-task condition. Using the same methods as described in the main text, age group difference in PFC activation was compared by a Wilcoxon’s rank sum test. The effects of age group and task conditions (single vs dual) in motor and cognitive performances were compared using aligned rank tests (3–5). Similar to the findings in the hard dual-task, older adults showed significantly higher PFC activation than younger adults during dual-task relative to rest standing. There was a significant interaction (trending in the hard dual-task) and a significant main effect of task (as observed in the hard dual-task) in the motor performance. In the cognitive performance, there was a main effect of age group like the hard dual-task, but there was also no main effect of task which was observed in the hard dual-task, suggesting that the cognitive performance change was only observable in the hard dual-task. This is not surprising considering the task difficulty was manipulated by increasing the cognitive task difficulty during the same motor task (walking).

| **PFC Activation** | | | | | | |
| --- | --- | --- | --- | --- | --- | --- |
|  | | **Older Adults**  **Mean (SD)** | **Young Adults**  **Mean (SD)** | **Test stats** | | |
| PFC activation during easy dual-task | | 3.60 ± 3.57 | -0.28 ± 1.55 | z = 3.98, p < 0.001 | | |
| **Performance** | | | | | | |
| **Domain** | **Task** | **Older Adults**  **Mean (SD)** | **Young Adults**  **Mean (SD)** | **Age Group Effect** | **Task Effect** | **Group x Task Interaction** |
| Motor  (speed, m/s) | Walk | 0.97 ± 0.13 | 1.01 ± 0.13 | F (1,40) = 1.51, p = 0.23 | F (1,40) = 106.80, p < 0.001 | F (1,40) = 5.36, p = 0.03 |
|  | Easy dual-task | 0.82 ± 0.14 | 0.92 ± 0.14 |  |  |  |
| Cognitive (rate, letters/s) | Standing cognitive task (Easy) | 0.50 ± 0.12 | 0.58 ± 0.10 | F (1,40) = 6.62, p = 0.01 | F (1,40) = 1.03, p = 0.32 | F (1,40) =1.66, p =0.21 |
|  | Easy dual-task | 0.51 ± 0.10 | 0.59 ± 0.10 |  |  |  |

Table S4. Neuropsychological evaluation result and Spearman’s correlations with locomotor savings. Yellow indicates trending relationships with 0.05 < p < 0.1. Reference refers to the age normed standardized score for a test. Abbreviated names of the cognitive tests were shown in the table with the full names included in the footnote (*). The trending relation observed between visuospatial index and locomotor savings is consistent with the literature often reporting associations between motor learning and visuospatial function (6–8).

| **Cognitive Domains / Test** | **Reference** | | **Mean** | **SD** | **Min** | **Max** | **rho** | **p** |
| --- | --- | --- | --- | --- | --- | --- | --- | --- |
| MoCA* | | / | 27.05 | 1.47 | 24.00 | 30.00 | 0.27 | 0.23 |
| RBANS† total index | | 100 ± 15 | 08.71 | 13.58 | 92.00 | 143.00 | 0.14 | 0.56 |
| Attention index | | 100 ± 15 | 111.14 | 13.44 | 91.00 | 138.00 | 0.09 | 0.71 |
| Visuospatial index | | 100 ± 15 | 105.43 | 15.98 | 81.00 | 131.00 | 0.40 | 0.07 |
| Immediate memory index | | 100 ± 15 | 103.43 | 13.53 | 81.00 | 140.00 | 0.04 | 0.88 |
| Delayed memory index | | 100 ± 15 | 105.81 | 10.50 | 84.00 | 127.00 | 0.14 | 0.56 |
| Language index | | 100 ± 15 | 104.00 | 6.83 | 92.00 | 120.00 | 0.11 | 0.64 |
| WRAT5‡ | | 100 ± 15 | 113.43 | 11.95 | 85.00 | 134.00 | 0.18 | 0.43 |
| Action verbal fluency | | 50 ± 10 | 48.95 | 10.13 | 37.00 | 74.00 | 0.04 | 0.86 |
| Executive function§ | | 10 ± 3 | 12.62 | 1.93 | 9.00 | 16.80 | 0.12 | 0.59 |
| Cognitive switching¶ | | 10 ± 3 | 11.83 | 2.79 | 4.50 | 16.50 | 0.06 | 0.81 |

*MoCA = Montreal Cognitive Assessment.

†RBANS = Repeatable Battery of the Assessment of Neuropsychological Status, which includes a total index score and five domain index scores as shown in the table (9).

‡WRAT5 = Wide Range Achievement Test Version 5.

§Executive function was measured with subtests of the Delis-Kaplan Executive Functioning System: Color Word Interference, Verbal Fluency, and Trail Making Test. A composite score was created by averaging scaled scores from Color-Word Interference condition 3, Color-Word test 4, Trail Making Test condition 4, and Verbal Fluency conditions 1, 2, and 3 total correct.

¶Cognitive switching was more specifically quantified by a composite score averaging Color-Word Interference test condition 3, Trail Making Test condition 4, and Verbal Fluency condition 3 category switching scaled scores from D-KEFS (10).

Table S5. Primary analysis of the correlation between locomtor savings and attentional control of walking repeated when using deoxygenated hemoglobin (Hbr) to quantify PFC activity. Following the recommendations of the fNIRS literature (11), all analysis results reported in the main text is reproducible using PFC activation quantified by Hbr as shown below. AGI is composed of PFC activity, which can be quantified by oxygenated hemoglobin (Hbo, shown in the main text) or Hbr shown here. Significant Spearman’s correlation was highlighted in red. Consistent with the findings reported in the main text and in Fig. S1, and Table S2, a positive relationship between attentional control of walking and locomotor savings was observed when using attentional gait index and this relationship exists in older adults only. Note that the relationship between locomotor savings and task performance change was not shown again because it remains the same regardless of which hemoglobin species was used to compute AGI.

|  | Older Adults | Younger Adults |
| --- | --- | --- |
| AGI computed from Hbr |  |  |
| Easy dual-task | $\rho=0.46, p=0.04$ | $\rho=0.34, p=0.14$ |
| Hard dual-task | $\rho=0.46, p=0.04$ | $\rho=0.21, p=$ 0.36 |
| AGI subcomponents (PFC activation measured by Hbr) | | |
| Easy dual-task | $\rho=-0.02, p=0.$94 | $\rho=0.13, p=0.57$ |
| Hard dual-task | $\rho=0.11, p=0.6$5 | $\rho=-0.12, p=0.$61 |

Table S6. Raw values and comparisons between age group in the attentional gait index computed from Hbr and PFC activation measured by Hbr. Wilcoxon’s rank sum tests were performed as in the main text. There was no significant age difference in either the attentional gait index or the PFC activation measured by Hbr. This is not surprising as Hbr has lower signal to noise ratio than Hbo (11).

|  | **Older Adults**  **Mean (SD)** | **Young Adults**  **Mean (SD)** | **Test Stats** |
| --- | --- | --- | --- |
| **AGI Computed from Hbr** | | | |
| Easy dual-task | 23.2 ± 10.82 | 21.14 ± 6.46 | z = 0.78, p = 0.44 |
| Hard dual-task | 44.84 ± 26.75 | 42.1 ± 26.19 | z = 0.18, p = 0.86 |
| **PFC activation measured by Hbr** | | | |
| Easy dual-task | 0.96 ± 4.33 | -0.63 ± 2.13 | z = 1.13, p = 0.26 |
| Hard dual-task | 1.02 ± -3.94 | -0.85 ± 2.52 | z = 1.21, p = 0.23 |
